# A geometric representation unveils learning dynamics in primate neurons

**DOI:** 10.1101/561670

**Authors:** Yarden Cohen, Elad Schneidman, Rony Paz

**Affiliations:** Dept. of Neurobiology, Weizmann Institute of Science, Rehovot, ISRAEL 76100

## Abstract

Primates can quickly and advantageously adopt new behaviors based on changing stimuli relationships. We studied acquisition of a classification task while recording single neurons in the dorsal-anterior-cingulate-cortex (dACC) and the Striatum. Monkeys performed trial-by-trial classification on a rich set of multi-cue patterns, allowing de-novo learning every few days. To examine neural dynamics during the learning itself, we represent each rule with a spanning set of the space formed by the stimuli features. Because neural preference can be expressed by feature combinations, we can track neural dynamics in geometrical terms in this space, allowing a compact description of neural trajectories by observing changes in either vector-magnitude and/or angle-to-rule. We find that a large fraction of cells in both regions follow the behavior during learning. Neurons in the dACC mainly rotate towards the policy, suggesting an increase in selectivity that approximates the rule; whereas in the Putamen we also find a prominent magnitude increase, suggesting strengthening of confidence. Additionally, magnitude increases in the striatum followed rotation in the dACC. Finally, the neural representation at the end of the session predicted next-day behavior. The use of this novel framework enables tracking of neural dynamics during learning and suggests differential yet complementing roles for these brain regions.

## Introduction

Learning to classify multi-cue stimuli in order to produce the correct action is an adaptive flexible behavior required from animals on a daily basis. Accordingly, such tasks are commonly used to explore learning strategies in humans (Gluck et al., 2002; Goodman et al., 2008; Lagnado et al., 2006; Nosofsky et al., 1992; Shepard et al., 1961), as well as clinical implications (Meeter et al., 2008; Shohamy et al., 2008; Speekenbrink et al., 2010; Stuss et al., 2000). Recent studies have shown that performance depends on complexity (Feldman, 2000) and can be predicted using models that rely on high-order features of the stimulus with individual priors (Cohen and Schneidman, 2013). Studies of rule-based classification in monkeys have ascribed a major and complementary role to the striatum and regions of the prefrontal-cortex (PFC) (Balleine et al., 2007; Seger and Miller, 2010). In paradigms that impose category boundaries on multiple stimuli, individual neurons in the PFC exhibit category preference to the different classes (Cromer et al., 2010; Freedman and Assad, 2016; Freedman et al., 2003; Gold and Shadlen, 2007; Histed et al., 2009; Kim and Shadlen, 1999; Muhammad et al., 2006; Wallis et al., 2001).

Within the PFC, the anterior-cingulate-cortex (dACC) widely projects to striatal regions (Averbeck et al., 2014; Heilbronner et al., 2016; Ongur and Price, 2000), and is involved in several cognitive functions that contribute to the learning process itself beyond the final representation of the rule. Neurons in the dACC represent attention, reflect actions that lead to reward, signal outcome of previous trials, and form and integrate representations of task structure (Chudasama et al., 2012; Haroush and Williams, 2015; Hayden and Platt, 2010; Heilbronner and Hayden, 2016; Kolling et al., 2016; Lee et al., 2007; Mansouri et al., 2009; Rudebeck et al., 2008; Rushworth and Behrens, 2008; Saez et al., 2015; Seo and Lee, 2007; Seo and Lee, 2009; Wallis and Kennerley, 2011). The striatum in turn, receives wide projections from the dACC and plays a role in choosing actions and supplies reinforcement signals which can help to establish a strategy during learning (Averbeck and Costa, 2017; Graybiel and Grafton, 2015; Jin and Costa, 2015; Kim and Hikosaka, 2013; Lau and Glimcher, 2008; Merchant et al., 1997; Seger, 2008; Seo et al., 2012; Williams and Eskandar, 2006).

Less is known about how single neurons form representations as learning progresses and gradually becomes relevant to the final classification being imposed. This is mainly because most studies follow extensive training and the neural correlates relate more to the final representation, perception, and recognition, and less to the gradual learning process. Classically, rule-based classification can take several forms(Seger and Miller, 2010), and seminal studies used few governing principles where the animals learn to assign different outcome probability or value (Padoa-Schioppa and Assad, 2006; Yang and Shadlen, 2007), acquire arbitrary stimulus-motor associations (Brasted and Wise, 2004; Buch et al., 2006; Mitz et al., 1991), or switch contingencies between the rules (Buckley et al., 2009; Wallis et al., 2001).

Here, we examine neural dynamics in the dACC and the striatum during learning of several de-novo classification tasks. To do so, we trained two monkeys (macaca fascicularis) to perform trial-by-trial classification learning based on visual patterns composed of ‘bits’ of black and white squares. Each session required to learn a classification rule on patterns of N=3 bits. Patterns were presented in a pseudo-random non-biasing order (Fig.1A, Methods). Out of the 256 (2^2^3) possible rules, we chose seven rules in which the correct label was determined according to either single, pairwise, or triple-wise dependencies between the bits in the pattern (Fig.1B). These rules are unbiased, namely equally partition the set of 8 patterns; and are independent of each other, namely learning a rule results in chance performance for all other rules. Additionally, we included the majority rule (also unbiased, Fig.1B). We repeated this set of eight rules after >4 weeks to obtain enough neurons recorded per rule. The monkeys did not experience any of them prior to recordings.

**Figure 1.**
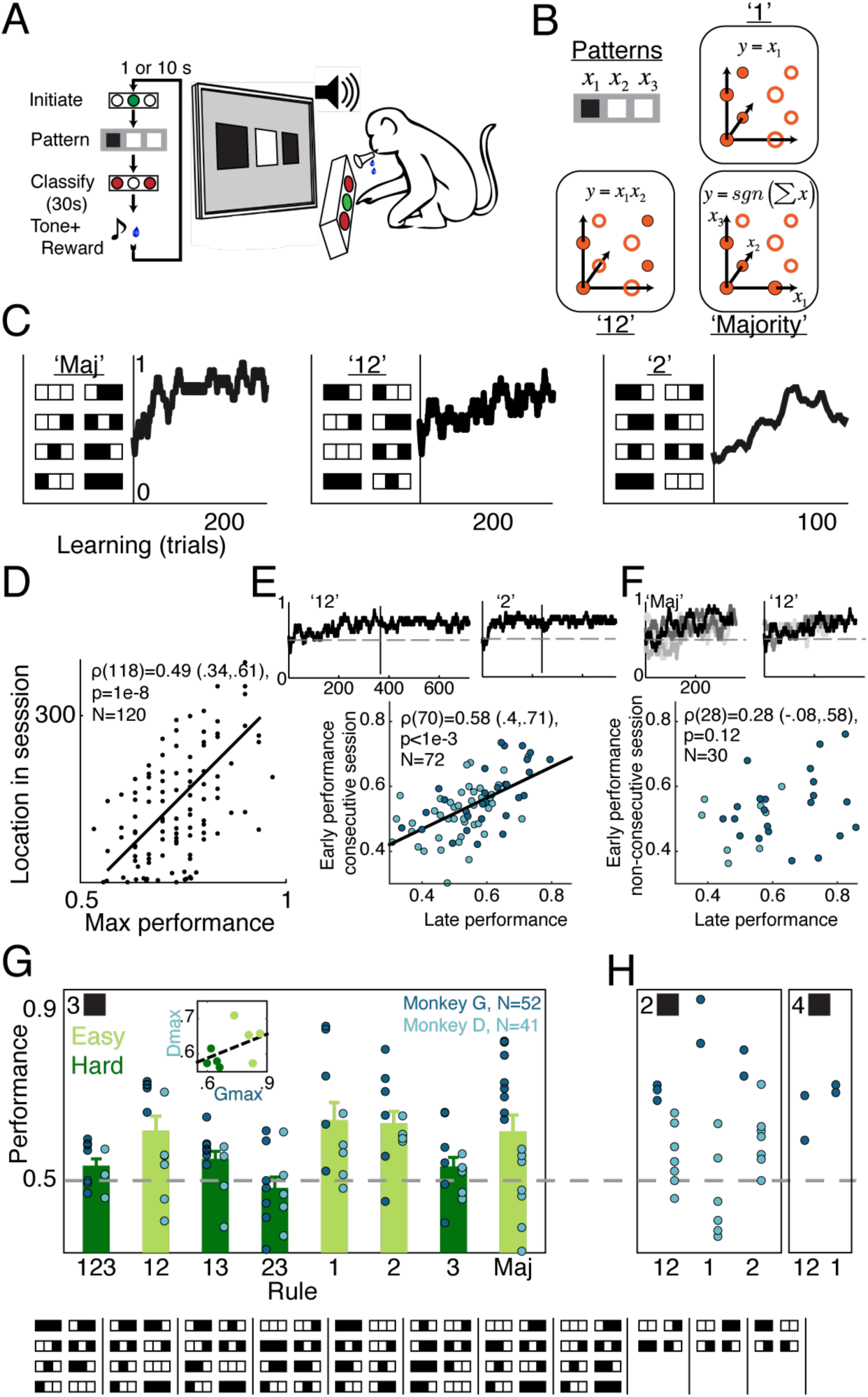
Monkeys learn classifications with varying complexity. **A**. Behavioral paradigm: pressing and holding the middle button initiates a new trial. After the pattern appears on the screen the monkey has 30s to classify it with the left or the right button. A fluid reward follows a correct choice and a short timeout after an incorrect choice. **B**. A scheme of the rule-based classification. Shown are: one-bit-rule ‘1’ (i.e. decision is based only on the identity of the first bit), two-bit-rule ‘12’ (decision is based on a XOR of bits ‘1’ and ‘2’), and the Majority-rule (decision is based on summing bits). In each panel the two categories are represented by full and empty circles in the space defined by the individual bits. **C**. Learning curves. Shown are individual learning curves from 3 example days each with a different rule (monkey G – two left examples, monkey D – right example). The curves are shown next to the underlying truth table of the rule where patterns are stacked by category. **D**. Continuous longer learning within-session is beneficial. Shown is the location within the session of the maximum performance plotted against the maximum performance (best 30 consecutive trials), for all sessions. **E**. Overnight retention. Shown are two examples of learning curves in consecutive days with vertical lines separating the sessions. Over all sessions, there was a significant correlation between performance at the end of a session and performance at the beginning of the subsequent session with the same rule (monkey D: light blue, r=0.48, p<0.001; monkey G: dark blue, r=0.61, p<0.001; Both: r=0.48, p<0.001). **F**. No retention between rule repetitions with >4 weeks separation. The use of eight different rules that use the same cues/stimuli over 4-5 weeks means it is virtually impossible to memorize the different rules. Shown are two examples of three sessions (scales of gray) that repeated the same rule but with >4 weeks between repetitions. Over all sessions, there was no relationship between performance at the end of learning a rule and performance at the beginning of next repetition of the same rule (monkey D: light blue, r=0.23, p>0.5; monkey G: dark blue, r=0.21, p>0.3; Both: r=0.12, p>0.2). **G**. Performance in all 3-bit rules, average performance in the last quarter of each session for all rules and all sessions, for both monkeys (D: light blue; G: dark blue). In addition, for each rule, shown is the mean and SE averaged over animals and sessions, and classified into ‘easy’ and ‘hard’ rules (in ‘easy’ rules performance is significantly above chance-level on average). Inset show maximal performance in each rule comparing the monkeys. Truth tables for all rules are shown below. **H**. Performance in 2-bit and in 4-bit rules (neural activity was recorded only during 3-bit rules).

## Results

### Learning classification rules

Both monkeys exhibited within-session learning (Fig.1C), continuous performance improvement in longer sessions (Fig.1D, Pearson r(118)=0.49(0.34,0.61), p<1e-8), as well as next-day retention (Fig.1E, Pearson r(70)=0.58(0.4,0.71), p<1e-3). As expected from learning eight different rules that change every few days but use similar visual cues, neither of the monkeys showed retention benefit over the month between rule repetitions (Fig.1F, Pearson r(28)=0.28(−0.08,0.58), p=0.12). Despite the hard task, both monkeys achieved learning of all rules in some sessions yet with different levels of accuracy that range across sessions and rules (Binomial tests, Fig.1G, Fig. S1), and with differences in overall performance as reflected also in learning of 2-bit and 4-bit rules (Fig.1H). These results are highly similar to the behavior of human subjects that exhibited similar behavioral variability across individuals, rules, and sessions(Cohen and Schneidman, 2013).

In pure stimulus-response associative learning, the correct response for each stimulus is acquired independently of other stimuli and independently of the general rule. Conversely, we found that pattern-specific error-rates following correct classification show strong dependence on the specific rule (Fig.S2, Kruskal-Wallis test,*χ*^2^(7,366) = 89, p < 1e-15), and interacted with performance on other patterns (Fig S3, Kruskal-Wallis test, *χ*(7,440)^2^ > 88, p < 1e-10). Moreover, even the stimulus-response learning of the more salient patterns showed variability across rules and strong dependence on the specific rule (Fig.S4, Kruskal-Wallis test, 24 < *χ*^2^(3,36) < 33, p < 1e-4). The rule-based learning was further supported by consistency across learning of different rules (Fig.S5), by the ability to learn 4-bit rules (Fig.1h, 16 patterns, 65536 possible rules, in one animal). Finally, we observed performance deterioration in pattern-specific associations following rule switch that do not require relearning those specific associations (Fig. S6), and that simple performance based strategy (win- stay, lose-switch) poorly describes answer sequences (Fig. S7, Wilcoxon’s signed rank test, p < 1e- 15). Overall, this evidence suggests that monkeys did not use only simple memorization of arbitrary stimulus-response associations, although it can still contribute to rule-learning here.

We conclude that both monkeys learned in a significant number of sessions with some rules being easier to learn than others. The complexity and richness lead to reasonable error rates and reflect a more natural success rate, compared to over-trained animals, and therefore allows comparison of neural responses from successful learning to unsuccessful trials, rules and sessions.

### Single neurons represent the classification rule

To examine and compare neural responses, we recorded single-units during learning of all 3-bit rules in the dACC (Brodmann area 24), Caudate and Putamen (Fig.2A,B, 543,115,114 neurons recorded in dACC, Caudate, and Putamen in 3-bit rule sessions). We divided the sessions into *easy* and *hard* rules based on the average success rate (Fig.1G). We first identified neurons that changed their post-stimulus spiking pattern to differentiate between the two category labels (*category-specific*, Fig.2C, left and middle), whereas other stimulus-responsive neurons maintained no category preference (*stimulus-specific*, Fig.2C, right). Overall, both the dACC and the Putamen had significantly more *category-specific* neurons during the late phase of sessions of *easy* rules compared to *hard* rules (Fig.2D, dACC: 21% vs. 12.5%, Pu: 22% vs 8% for *easy* vs. *hard*, binomial comparison z-test, dACC: z=2.43, p<0.01, Pu: z=2.12, p<0.02). The same was obtained when dividing sessions into high and low performance independent of the specific rule, namely session-based performance rather than rule-based performance (Fig. S8, dACC: z=2.046, p<0.021, Pu: z=1.79, p<0.04), and this was the case in each monkey separately (Fig.S8). Moreover, there was a monotonic relation between instantaneous performance and the proportion of neurons showing correct classification (Fig.2E).

**Figure 2.**
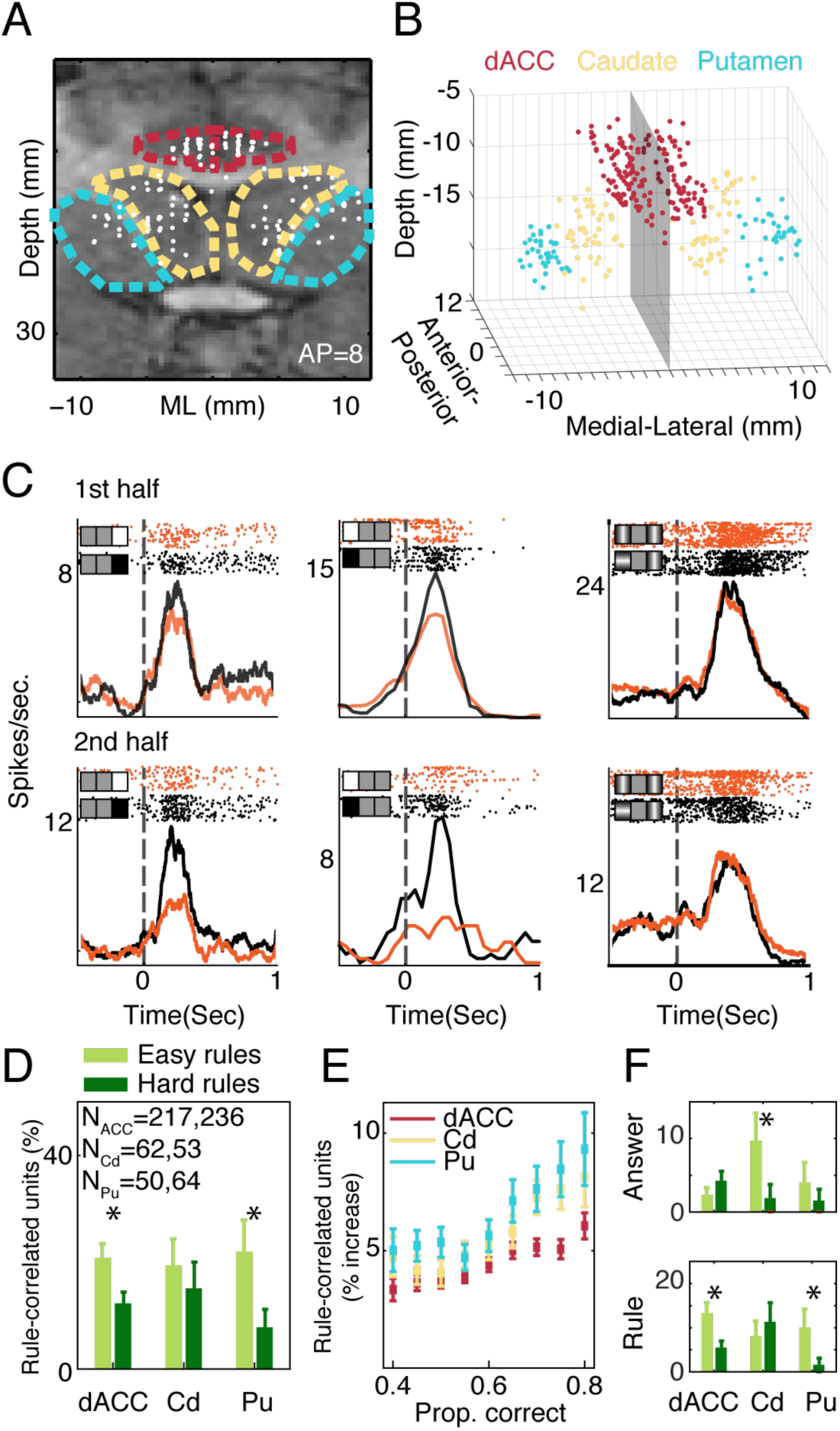
Neurons represent learning of classification. **A**. Recording locations projected on a coronal MRI section. Recording locations span ∼12mm in the anterior-posterior axis (hence some locations seem to be outside the regions of interest). Red marks borderline of the dACC, Yellow for Caudate and cyan for Putamen. **B**. 3-D reconstruction of all recording locations. Gray plane is the midline, anterior-posterior zero is at the anterior-commissure and depth is measured from the dura surface. **C**. Rule-learning in neurons. Spiking patterns are divided by the category label (orange and black rasters and PSTHs). Spike times are aligned to the stimulus onset (dashed line). The order of trials is top to bottom. Trials are divided into the first half (upper row) and second half (bottom row) of the sessions. The two left neurons differentiated between categories in the second half significantly more than in the first half, whereas the right neuron did not learn to differentiate categories. **D**. Fraction of single-units whose firing correlated with the rule at the end of learning (Mean and SE). Sessions are divided according to easy vs. hard rules, with more neurons signaling correct classification in easy rules in the dACC and the Putamen (p<0.05 for both, binomial tests). **E**. Instantaneous performance parallels neural-based categorization. The fraction of neural segments with significant category correlation increases with the mean performance (calculated by rolling regression windows of 40 trials in steps of 4 trials). **F**. Proportion of neurons with stronger correlation to the rule (bottom) and with stronger correlation to the actual choice (top), over both successful and error trials (William’s test), and comparing easy vs. hard rules. The dACC and the Putamen are significantly correlated more with the rule and not with the actual choice, whereas caudate neurons are significantly correlated with the choice only and not with the rule (significant comparisons are p<0.05, binomial tests).

Since the category label and the actual choice are correlated during correct performance, we used the occurrence of error trials to distinguish them and compared the correlation between neurons’ activity and the correct category to the correlation with the actual choice over trials (Fig.2F, William’s test). In sessions of *easy* rules, more neurons developed category correlations in the dACC and Putamen, whereas significantly more neurons developed actual-choice correlations in the Caudate (binomial comparison z-test, dACC: z = 2.88, p<0.003, Cd: z=1.72, p<0.05, Pu: z=2, p<0.03). Finally, in accordance with the aforementioned behavioral findings (Fig.S1-7), even under lenient conditions, only very few neurons (3%, p>0.1, Binomial tests) exhibited activity that can be attributed to memorizing of a specific stimulus-response, and moreover, most of these neurons were selective and sensory-specific to the all-black or all-white patterns (36/40, Fig. S9).

Together, these findings suggest that the modulation of single neuron activity in the recorded regions reflect different roles during learning. Whereas the dACC and Putamen reflect more the learned classification, the Caudate reflect more the actual choice. We therefore focus in the following analyses on the dACC and the Putamen.

### A geometric representation to track neural dynamics during learning

We next examine how the modulation of neural activity occurs dynamically and continuously during the learning process itself. To track representations during acquisition, we note that each pattern can be represented by its visual features (Fig.3A), and if we choose a complete set of statistically independent binary functions as features (a minimal spanning basis), we can describe any rule as a weighted combination of these features. In other words, each rule is a vector in this feature space. This choice of the basis also guarantees that a neuron’s selectivity is defined by the correlation between its activity and the features (*neural-vector*). In turn, this allows the use of rolling regression to identify the neuron’s dynamic preference, depicted by the trajectory of its *neural-vector* in this space (Fig. S10, Fig.3B).

**Figure 3.**
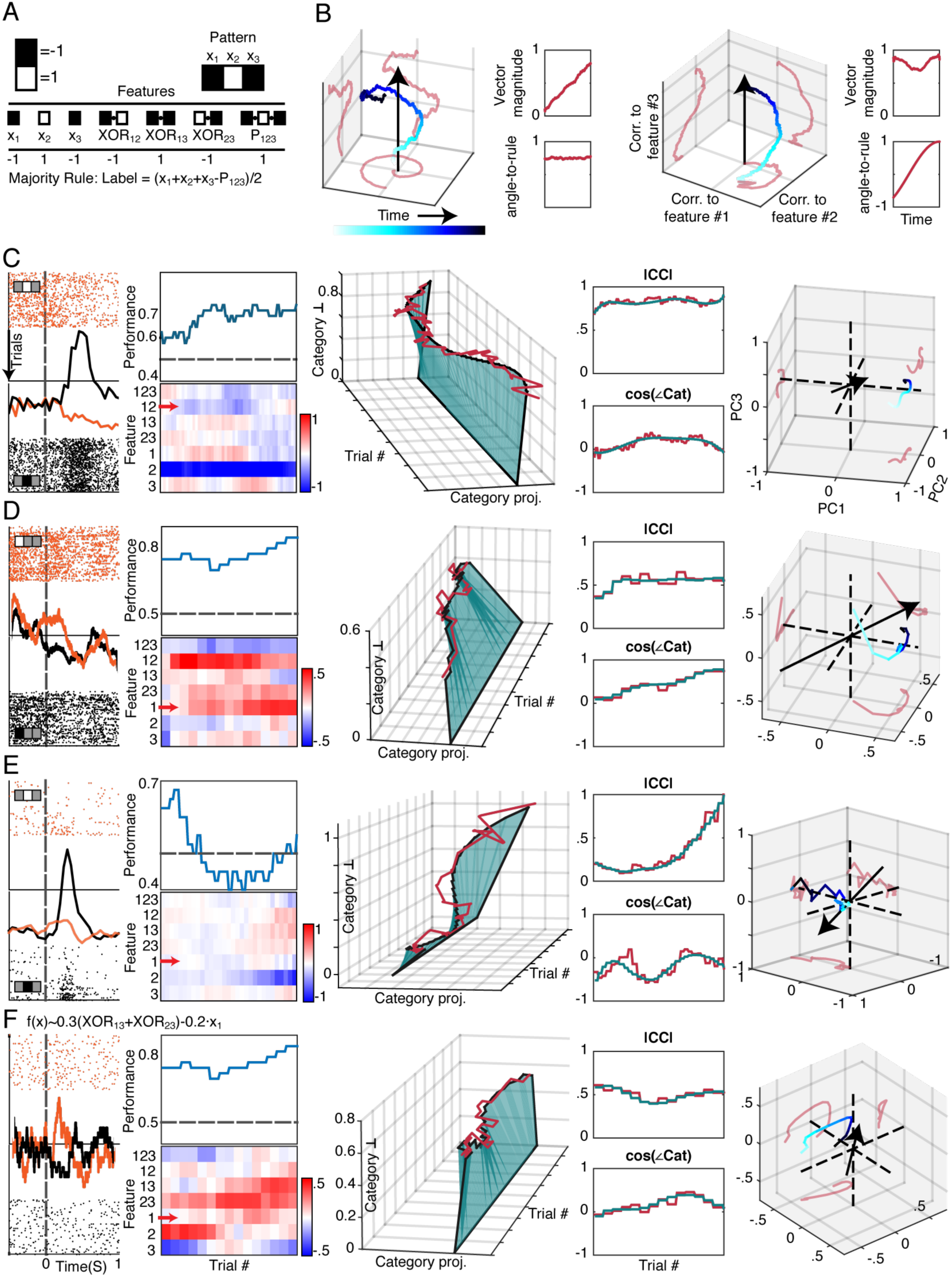
A geometrical representation reveals single-neuron dynamics during learning. **A**. The 7 features that form a linear basis (minimal and spanning) for the space in which all 3-bit rules reside. Shown is an example for the representation of the pattern 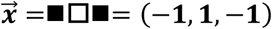 by its parity features of: first order, 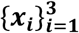, second order, 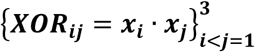, and third order, ***P*_123_** = ***x***_**1**_ · ***x***_**2**_ · ***x***_**3**_ and the representation of the majority rule as a combination of these features. **B**. A schematic demonstrating that in this feature space, the change in a neuron’s response modulate its projection on the vector that represents the rule (the black arrow). An increase in the projection can result from two processes: a trajectory change that increases the neural-vector magnitude (left) or neural-vector rotation towards the direction of the rule-vector (right), or of course both. The trajectory is color coded for time and red curves depict its projections on the axes. Insets separate the dynamics of vector-magnitude (top) and angle-to-rule (bottom). For presentation purposes only, the neural trajectory is plotted in the space of the three main principle axes (PCA), yet for all analyses the neural-vectors were computed in the full 7D space. **C-F**. Examples of single-neuron neural-vector dynamics. Each row is a neuron and shows, from left to right:

1. Rasters aligned to stimulus onset with trials advancing from top to bottom and divided by the sign of the preferred feature at the second half of the session. PSTHs for the top raster (orange) and for the bottom one (black).
2. Top: the behavioral learning curve (20 trials running average). Bottom: the regression correlation coefficients (blue-red color-bar) for each feature in the basis (y-axis) along all trials (x-axis). Red arrow marks the feature that defined the correct label in the specific session.
3. The norm projections of correlation vectors (red) on the category (x-axis) and its orthogonal subspace (Category ⊥, z-axis) over time (y-axis). The blue surface connects (0,0) to the smoothed dynamics to visualize rotation and extension.
4. The correlations’ vector magnitude (top) and angle to category (bottom) plotted over time. Red: raw data. Blue: smoothed data.
5. The high dimensional (7D) neural trajectory projected on the three main principle axes (PCA) and curve fitted, color-coded for time progress in the session (light → dark blue). The black arrow is the rule vector in that session and the red curves are the projections of the trajectory on the axes (principal components). **C**. A neuron that does not change its selectivity during learning **D**. A neuron that rotates towards the correct category by learning the right feature **E**. A neuron that extends its vector towards a ‘wrong’ feature. **F**. A neuron that exhibits a complex relationship of vector extension and rotation.

Importantly, because the *neural-vector* is described in terms of movement in features’ space and its relation to the *rule-vector* that varies in different sessions, this geometric framework allows comparing neural dynamics across sessions of different rules. Specifically, the projection of the *neural-vector* onto the *rule-vector* is equivalent to the correlation between neural activity and the categories determined by the rule (see methods). To increase its category representation, a neuron can employ two distinct processes that contribute to a higher agreement (projection) between the *neural-vector* and the *rule-vector*: increasing vector magnitude (Fig.3B, left) – reflecting a confidence strengthening (being ‘louder’), or by rotating towards the direction of the *rule-vector* (Fig.3B, right) - reflecting a policy change. We note that choosing any other representation, by another feature space, can be expressed as a weighted combination of the feature basis, and so our findings do not depend on the specific set of features we used.

Representing the high-dimensional trajectories in the feature-space (7-d) by changes in angle-to-rule and vector-magnitude reveals several types of single-neuron dynamics. Whereas some neurons remained selective to a feature regardless of the performance improvement (Fig.3C), other neurons rotated towards the correct rule during learning (Fig.3D. Fig. S11), and yet other neurons increased their vector magnitude and developed feature selectivity but with no relationship to the correct rule (Fig.3E). Interestingly but somewhat expected, many neurons performed a more complex path and their trajectories involved changes in several features in parallel (rotation and magnitude, Fig.3F, Fig. S12). The geometric representation allows interpretation for these complex paths as well – such as rotation in the orthogonal subspace (‘Category ⊥’, e.g. Fig.3F), and magnitude increases in building preference to a combination of several features, that eventually leads to significant agreement with the *rule-vector* (see Fig.S12 for further examples).

### Neural trajectories closely match behavior

Next, we asked if this neural representation agrees with the behavioral changes in individual sessions. We computed the correlation between the behavioral performance and the *neural-vector*, separately for angle and magnitude. There was a large and significant proportion of neurons in both regions for which changes in angle-to-rule were correlated with the behavior (Fig.4A, 32% of the dACC neurons and 37% of the Putamen neurons, Pearson, p<0.01). Similarly, a significant proportion of neurons changed their vector-magnitude in correlation with performance (Fig.4B, 19% of the dACC neurons and 18% of the Putamen neurons, Pearson, p<0.01). These populations intersect such that many neurons with vector-magnitude changes also exhibit angle-to-rule changes (Fig. S13).

**Figure 4.**
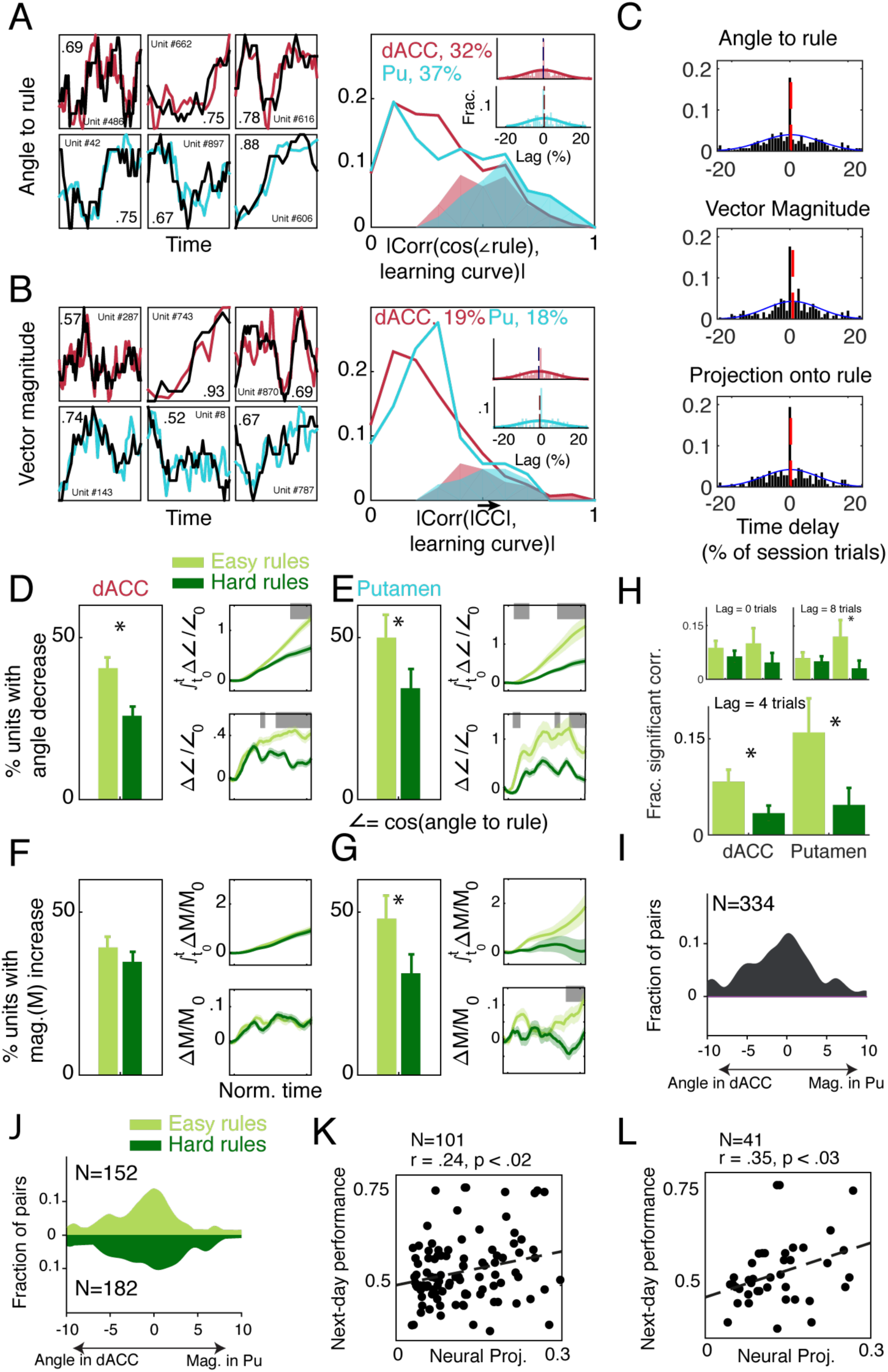
Neural dynamics match learning behavior and varies across regions. **A**. Change in angle-to-rule (**cos**(**∠*Rule***)) during a session overlaid with the performance behavior (black), showing a highly similar temporal pattern. Three dACC neurons (upper row, red) and three Putamen neurons (lower row, cyan), all with significant correlation between neural dynamics and performance (p<0.01 for all, Pearson). Right panels show histograms of correlation coefficients for all neurons, with shaded area mark neurons exhibiting significant correlations at p<0.01. Both regions contained a highly significant number of neurons with dynamics similar to behavior (p<0.01 for both, *χ*^2^). Insets show the distribution of trial lags between neural dynamics and behavior, with the mean lag not different than zero (p>0.1, t-tests). **B**. Same as in (a) for dynamics in vector magnitude (|*CC*|). **C**. The distribution of trial lags between neural dynamics of dACC neurons and of Putamen neurons, for all simultaneously recorded pairs in both regions (p>0.1 for all, t-tests). Shown are the neural lags when taking into account only angle (upper), magnitude (middle), and combined projection on the rule-vector (bottom). **D**,**E**. Proportion of neurons that significantly decreased their angle-to-rule during the session was different in easy rules vs. hard rules in both the dACC (**d**) and the Putamen (**e**) (p<0.05 in both, binomial z-test). Right panels show the population-average angle change (bottom, SE in shaded color) and the normalized cumulative change (top). Sessions were time-warped for averaging. Gray bars indicate significant difference between easy and hard rules (p<0.05, bootstrap). **F**,**G**. Same as in (d,e) for change in vector magnitude. However, in contrast to rotation in the both regions, only the Putamen showed significant increase in the vector-magnitude in easy vs. hard rules. **H**. Local shifts in angle-to-rule follow the behavior in more neurons when comparing easy to hard rules in both regions, but only in a lag of 4 trials (main panel) and not in zero or 8 trials lag (top insets). **I**. Distribution of optimal lags for all simultaneously recorded pairs of neurons, between change in vector magnitude of the putamen neuron and angle-to-rule of the dACC neuron. The mean is significantly below zero, indicating that changes in angle in dACC neurons preceded changes in magnitude in Putamen neurons. All other combinations were not significant (Fig.S8). **J**. Same as in (I) but separately for easy and hard rules. **K**. Neural projection at the end of a session predicts next day performance. The projection of each Putamen neuron’s activity at the end of the session (neural-vector) onto the next-day rule (rule- vector), against the mean performance in the beginning of the next-day session (the neural-vector is averaged over the last 25% of the session and performance is averaged over the first 25% of the next day session), showing a positive correlation (r=0.24, p<0.02, black regression line). **L**. Similar to (K) but when the rule changed overnight, and the neural-vector at the end of the session is projected onto the next-day rule, showing a positive correlation (r=0.35, p<0.03)

We tested for the possibility of a temporal lag between the neural dynamics and the behavior. In all cases, for both regions and for both magnitude and angle changes, neurons were equally distributed between preceding change in behavior and slightly following it (Fig.4A,B, right-insets, means not different than zero, p>0.1 for all, Wilcoxon’s signed rank tests). Importantly, because neurons were recorded simultaneously in both regions, we could test if there is a lag in canges of either angle or magnitude between pairs of neurons recorded from the dACC and Putamen within the same session. This revealed a wide distribution (Fig.4C), but without a specific directional lag (p>0.1 for all, Wilcoxon’s signed rank tests), suggesting that information is being shared in both directions with both short lags (within the same trial) but also longer ones (over few trials). This is in close agreement with previous findings of information transfer in corticostriatal loops during learning.

### Representations of confidence and policy across regions

The similar temporal dynamics between neural trajectories and behavior allows relating the neural representations to the different components of learning: increasing vector magnitude reflecting a confidence increase, and rotating the vector towards the rule reflecting a policy change. We find that in *easy* rules, more neurons rotated towards the rule (diminished their angle-to-rule), in both the dACC and the Putamen (Fig.4D,E, binomial z-test comparing *easy* to *hard* rules, dACC: 40% vs 25.5%, Putamen: 49% vs 34%, dACC: z=3.32, p<0.001, Pu: z=1.68, p<0.05). This rotation gradually progressed along the session as performance improved (Fig.4D,E– right panels, Fig. S14).

However, a qualitative difference between the dACC and Putamen is revealed when examining vector-magnitude changes. In the dACC, there was no difference in the number of neurons with vector-magnitude changes between *easy* and *hard* rules (Fig.4F, binomial z-test. dACC: 38% vs 34%, p>0.1). In contrast, in the Putamen, significantly more neurons increased their vector-magnitude in *easy* rules (Fig.4G, binomial z-test. Putamen: 47% vs 31%,z=1.82, p<0.04). These changes became much more prominent towards the end of the session, suggesting strengthening (confidence) of the policy that developed earlier (compare Fig.4D,E, right-panels to Fig.4G, right-panels; results remain similar when re-labelling the *Majority* rule as *easy* for monkey-G and *hard* for Monkey-D, since it is the only rule on which the monkeys’ behavior did not completely agree, Fig. S15).

In contrast to the angle and magnitude, both the dACC and Putamen similarly represented the actual-choice in *easy* compared to *hard* rules (Fig.S16E,F), reinforcing the above findings that the dynamics represent learning. As validation and in agreement with Fig. 2F, Caudate neurons represented the actual-choice differentially in *easy* rules (Fig.S16D), but not for angle-to-rule and vector-magnitude (Fig.S16A-C)

Together, these findings suggest that both the Putamen and dACC reflect a process of policy search for the correct rule, but mainly the Putamen reflects strengthening and confidence gain that accompanies successful learning.

### Differential dynamics across regions

To examine more closely if the changes in neural properties relate to behavioral performance, we examined the correlation between the learning curve and the change in angle-to-rule, namely the change in subsequent windows (rather than the overall angle-to-rule as in Fig.4A). We then compared the proportion of neurons that showed significant correlations in *easy* vs. *hard* rules and found that more neurons showed a change in angle-to-rule that followed behavioral changes in *easy* than in *hard* rules. This was so in both the striatum and the dACC when we computed the correlation with a lag of 4 trials (Fig.4H, binomial z-test. dACC: z=2.24, p<0.02, Pu: z=2.03, p<0.03), and diminished when the lag was higher (8 trials, Fig.4H top-right) or with zero lag (Fig.4H top-left). There was no difference when the lag was negative, namely when neural change precedes behavioral change. This suggests that when a search for answers succeeds or fails beyond average, it is followed by a *neural -vector* rotation towards the rule. The fact that the rotation was directed towards the daily rule indicates that it is not reward alone but instead due to success/failure.

We exploited the simultaneous recordings to test if the two properties, magnitude and rotation, have differential dynamics between regions. We took the vector-magnitude and the rotation-to-rule for each neuron and computed the optimal lag (as in Fig.4C) between the two properties for all simultaneously recorded neurons. Only the lags between vector-magnitude in Putamen and angle-to-rule in dACC was biased, with the Putamen magnitude following dACC rotation (Fig.4I, Wilcoxon’s signed-rank test, z=-2.81, p<0.005; mean lag −2.8±0.96 trials; Fig.S17, all other comparisons were not different than zero, p>0.1 for all, Wilcoxon’s signed-rank tests). This was driven mainly by neural pairs recorded during *easy* rules (Fig.4J, Wilcoxon’s signed-rank test, z=-2.64, p<0.01, Mean lag - 3.56±1.36 trials).

Together, the results suggest that once a strip of successful responses occur, the *neural-vector* in both regions rotate towards the rule, and the rotation in the dACC is followed by magnitude extension in the striatum, likely strengthening the new policy.

### Neural vectors predict next-day (overnight) behavior

If the neural trajectories indeed reflect learning and changes in behavioral policy during a session, then the *neural-vector* at the end of a session represents the acquired policy and might predict early performance in the next-day daily session (a putative overnight retention process).

To examine this, we projected each neuron’s *neural-vector* at the end of a session onto the *rule- vector* of the following day, and computed the correlation between these projections and the early next-day’s performance. In Putamen neurons, *neural-vectors* predicted next-day success rates (Fig.4K, Pearson’s r(99) = 0.24 (0.05,0.42), p<0.015; (as control, we reversed the days and found no correlation). Namely, the closer the *neural-vector* at the end of the day to the next-day rule, the better is the initial performance.

We further tested if these neural priors could also bias learning of new rules in sessions when the next-day rule change. We repeated this analysis only for pairs of days when the rule changed in the second day, and again found a significant correlation for Putamen neurons (Fig.4L, Pearson’s r(39) = 0.35 (0.05,0.59), p<0.025; no correlation when reversing days as control). In other words, the closer the *neural-vector* at the end of a day to the *rule-vector* of the next day, the better will be the animal initial performance.

Together, these findings further strengthen the link between the neural geometric representation and the behavior, establish that the neural projection is a valid representation for the learned rule, and further demonstrate that it can bias future behavior.

## Discussion

We recorded and analyzed dynamic changes in neural representation while monkeys learned to classify multi-cue patterns based on a rich set of potential rules. In these natural conditions of varying eight different rules, the animals’ performance varied across sessions and rule types as previously shown in humans (Cohen and Schneidman, 2013). Nevertheless, we find that neurons in both the striatum and the dACC change their firing pattern to eventually represent the learned rule. To address the main challenge of characterizing single-unit dynamics during de-novo learning of rules, we developed an approach that represents each rule in the space formed by a set of features. Because this set of features is a minimal spanning basis, other cartesian coordinates that allow a vector representation are equivalent to it, making the findings generalizable and independent of the specific chosen features. We could therefore represent each neuron activity by two universal traits – its angle to the rule direction and its magnitude. These two measures have a natural interpretation, the rotating angle reflects learning-related policy or strategy changes, and the magnitude increase reflect strengthening or neural confidence (we do not claim a direct link to cognitive confidence). We suggest this has the benefit of denoting via these two geometric traits all rules that can be formalized by any high-dimensional stimuli space, and hence compare the dynamics of learning across tasks and rules.

Using this framework, we found that in both regions there is a large number of neurons that were dynamically synchronized with the behavior, either in their angle or in their magnitude. This suggests that this neural representation indeed captures the learning process (Brasted and Wise, 2004) and reorganizes to adapt to the new conditions (Golub et al., 2018). The representation revealed a dissociation in functionality: neurons in the dACC rotated to decrease their angle-to-rule, namely changed their strategy; whereas neurons in the Putamen changed their activity to reflect both strategy and confidence, by magnitude-increase of the neural-vector that likely reflects strengthening and reinforcement of the correct strategy (Graybiel and Grafton, 2015). In line with this interpretation and the role of the striatum in reinforcement, we find that rotation changes in the dACC were followed by magnitude extensions in the striatum, and even more so in successful learning.

In addition, apart from representations of the learned rule, we could also identify neurons with stable or changing correlations in the subspace of the features (Chen et al., 2001; Genovesio et al., 2005; Sadtler et al., 2014). Interestingly, such changes were observed during both successful and unsuccessful learning. Although the exact role of these neurons is yet to be determined, they likely reflect a trial-and-error search or retention of previously learned rules. This means that even if some tasks are not learned successfully, some neurons still approximate the rule during the learning, yet at the end the overall population does not.

In contrast to the Putamen and the dACC, Caudate neurons showed stronger reflection of the monkeys’ instantaneous actual-choice. Because in visuomotor associations the dorsal striatum represents actions’ value (Lau and Glimcher, 2008) and follows the choice representation in lateral PFC (Seo et al., 2012), yet the role of lateral PFC in category learning and generalization is still under debate (Minamimoto et al., 2010); our observations can support the notion that in rule learning Caudate activity reflects value-based action selection (Desrochers et al., 2015; Kim and Hikosaka, 2013; Williams and Eskandar, 2006; Yanike and Ferrera, 2014). Finally, neurons in the Putamen were related to overnight retention, and their representation at the end of a day predicted next-day behavior. Because we imposed a complex-hard task and learning usually continued over few sessions, this demonstrates that the striatum represents the policy that is used for retention, either within the striatum or by transfer to other regions (e.g. as in consolidation processes). This suggestion is in line with studies showing that the striatum maintains intermediate representations, potentially via sustained activity (Deffains et al., 2016), to allow learning that combines reinforcement and memory under spaced conditions(Doll et al., 2015; Wimmer et al., 2018).

Our results can further suggest how abnormalities in the cingulate-striatal network can result in maladaptive learning processes that lead to applying incorrect rules, and in extreme cases lead to psychopathologies(Averbeck and Chafee, 2016; Hyman et al., 2006; Lee, 2013; Salzman and Fusi, 2010). Overall, we present here a new computational framework to examine dynamics of neural changes, and suggest complementing roles for the dACC and Striatum in learning and retention of classification rules.

## Acknowledgments

We thank Yossi Shohat for animal training, welfare and experimental procedures; Dr. Yoav Kfir for scientific consult, Dr. Eilat Kahana for help with medical and surgical procedures; Dr. Edna Furman-Haran and Fanny Attar for MRI procedures. E.S. was supported by a European Research Council Grant 311238, an Israel Science Foundation Grant 1629/12, research support from Martin Kushner Schnur and Mr. and Mrs. Lawrence Feis, and a CRCNS grant. R.P. was supported by ISF #2352/19 and ERC-2016-CoG #724910 grant.

## Author contributions

Y.C, E.S. and R.P. conceived and designed the study; Y.C. performed the experiments and analyses; Y.C., E.S. and R.P. wrote the manuscript.

## Methods

All surgical and experimental procedures were approved and conducted in accordance with the regulations of the Weizmann Institute Animal Care and Use Committee, following National Institutes of Health regulations and with accreditation from the Association for Assessment and Accreditation of Laboratory Animal Care International.

### Animal training

Two male monkeys (Monkeys G and D, macaca fascicularis, 4-6kg) participated in the experiment. Before data collection, each monkey went through a training phase that acquainted it with all the task components and their sequence in a learning session. Both monkeys were trained similarly. They first learned rules with 2-bit patterns to understand the concept of the task. Then, immediately before the surgery, they experienced three rules with 3-bit patterns so that they would not be surprised when they see 3-bits for the first time during recordings. Importantly, these were 3 different rules than the 8 rules tested during recordings and shown in the manuscript (there are 256 possible for assigning 8 patterns into 2 categories). Other than that, no training was done. Therefore, in the recording sessions the monkeys were familiar with the concept of patterns and classification (Fig.1A). However, the monkeys did not experience any of the rules reported in this manuscript before the electrophysiological recordings began.

### Experiment sessions

The monkeys learned to classify binary patterns of N=3 squares. In each session, the entire set of 2^N^ possible patterns was presented. The order of patterns was generated by concatenating full sets of randomly ordered 2^N^ patterns. This process ensured that all patterns appear with the same temporal frequency and that no choice of behavioral rule, apart from the correct one, is beneficial in large portions of the session. For compactness we refer to the rules by their constituent squares. So, for example, in rule ‘3’ the label is determined by the color of the 3^rd^ square and in rule ‘12’ the label is determined by the XOR of squares 1 and 2. See Fig.1 and main text for the list of rules used in this study and during recordings. Each classification rule was replaced every few days and repeated after about a month and after a full cycle of the 8 rules was presented.

### Neural recordings

A craniotomy was performed under deep anesthesia and aseptic conditions and a recording chamber (27×27mm) was implanted above the midline and anterior commissure to allow daily electrodes insertion. The chamber’s positioning was done according to MRI calculated coordinates with respect to the identified bone structure around the ear canals and eye sockets. Still images were taken during the surgery to record the location of the chamber, the head holder and the screws on the skull for easier extraction process.

After surgery the monkeys were treated with analgesics (Buprenorphine) and antibiotics (Rocephin, Baytril). The monkeys were allowed to recover for 1-2 weeks before the first head restraining in the setup. The fluid consumption regime was gradually reinstated starting two weeks after surgery.

### MRI-Based Electrode Positioning

Anatomical MRI scans were acquired before, during, and after the recording period. Images were acquired on a 3-Tesla MRI scanner: (MAGNETOM Trio, Siemens) with a CP knee coil (Siemens). A T1-weighted, three-dimensional gradient-echo (MPRAGE) pulse sequence was acquired with a repetition time of 2,500 ms, an inversion time of 1,100 ms, an echo time of 3.36 ms, an 8 flip angle, and two averages. Images were acquired in the sagittal plane, 192 × 192 matrix, and 0.63 mm resolution. The first scan was performed before surgery and used to align and refine anatomical maps for each individual animal (relative location of the dACC and the Striatum, and anatomical markers such as the interaural line and the anterior commissure; confirmed using atlas). We used this scan to guide the positioning of the chamber on the skull at the surgery. After surgery, we performed another scan with 2-4 electrodes directed toward the dACC, Putamen and caudate. The regions’ depth was calculated from the dura surface and the plane of the top of the chamber. We assessed estimation of electrode tip locations and comparison to the MRI image with <1mm accuracy (mean=0.5mm).

### Mapping recording regions

During the first week of electrode insertions we performed a mapping procedure to identify the depth of cell bodies in prominent recording regions. During that week no behavior recordings were made and the fluid restriction was gradually reinstated.

Additionally, with every electrode insertion during the experiment we recorded the depths of cell bodies and were able to reconstruct the boundaries of our regions of interest.

### Electrophysiology

The monkeys were seated in a dark room and each day, up to six microelectrodes (0.6–1.2 M**Ω** glass coated tungsten, Alpha Omega) were lowered inside a metal guide (Gauge 25xxtw, outer diameter: 0.51 mm, inner diameter: 0.41 mm, Cadence) into the brain using a head-tower and electrode-positioning-system (Alpha-Omega). The guide was lowered to penetrate and cross the dura and stopped once in the superficial layer of the cortex. The electrodes were then moved independently further into either the dACC, Caudate or Putamen. Electrode signals were pre-amplified, 0.3 Hz–6 kHz band-pass filtered, and sampled at 44 kHz; and online spike sorting was performed using a template-based algorithm (Alpha Lab SNR, Alpha Omega). We allowed 15-30 minutes for the tissue and signal to stabilize before starting acquisition and behavioral protocol. At the end of the recording period, offline spike sorting was further performed for all sessions to improve unit isolation (offline sorter, Plexon).

### Data Analysis – Behavior

#### Performance

Each learning session results in a series of correct and incorrect answers, {*y*_*t*_}_*t*=1:*T*_ ∈ {0,1}^*T*^, T being the number of trials. To measure learning behavior and account for erratic tendencies we took the following steps:

1. To avoid the behavioral decline that may bias performance at the end of the sessions we disregarded up to the last 10% of the session if it contained only wrong answers. On average we ended up ignoring ∼1% or 2-3 trials in each session.
2. We define performance at the end of the session by averaging correct and incorrect answers in the last quarter of the session, 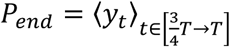. The confidence level for rejecting the null hypothesis of chance performance follows the regularized incomplete beta function, 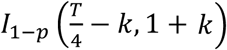, where 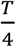 is the number of trials in the last quarter of the session and *k* is the number of correct answers during that segment (Fig. S1).
3. Identically to *P*_*end*_, we define *P*_*start*_ as the mean performance during the first quarter of the session.
4. In Fig. 1D, we define maximal performance as the best mean performance in 30 consecutive trials, 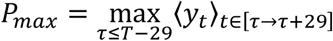.
5. In Fig.S2, Fig. S3, Fig. S4, and Fig. S6 we use pattern-specific performance. The order of pattern presentation, randomized batches containing all 2^N patterns, guaranteed that the sequences of pattern-specific presentations were perfectly-interleaved – allowing for the comparison of pattern-specific errors conditions on prior presentations of the same pattern (Fig. S2) or other patterns (Fig. S3). Similarly, the comparisons of pattern-specific learning curves is temporally-aligned between rules (Fig. S4) and with the general performance (Fig. S6).

#### Easy and hard rules

We label rules according to the monkeys’ ability or inability to recurrently achieve high performance in learning those rules. Thus, given that a subset of rules is labeled ‘easy’ and another subset is labeled ‘hard’, we computed the amount of variance, within the set of *P*_*end*_’s that the labeling explains. The R^2^ value is: 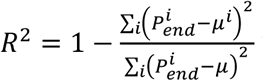 where, *μ* = ⟨*Pend*⟩ is the mean end-performance of all the sessions and *μ*^*i*^ are the mean end performances of sessions of either ‘easy’ or ‘hard’ rules (Fig.1G). We also labeled individual sessions as ‘high’ or ‘low’ performance independent of the rules to make sure our results are robust (Fig.S8).

#### Rule repetition effects

Each classification rule was used for 1-5 consecutive days and repeated after about a month. We compare the monkeys’ performance at the end of a session (*P*_*end*_(*n*) with ‘n’ standing for the n’th session of the rule) to the mean performance in the first quarter of the following session of the same rule, *P*_*start*_(*n* + 1). The comparison is made by calculating the Pearson correlation between the *P*_*end*_’s and *P*_*start*_’s.

We examined two distinct cases:

1. Taking sessions only from consecutive days, we calculate the correlation, *ρ*_*across*_, of the across-days learning (Fig. 1E).
2. Taking only sessions from the end of a consecutive sequence and the beginning of the following sequence, we calculate the correlation, *ρ*_*recall*_, of the monkeys’ ability to recall rules they encountered a month before (Fig. 1F).

#### Testing for pattern-specific memorization

We first consider a memorization scheme in which subjects perfectly learn a list of correct pattern-label pairs but do not generalize. The acquisition of such memorized patterns can be an all-or-none event, which means that after a certain pattern-label pair was memorized it will dictate choice behavior. Alternatively, we consider a gradual probabilistic association strengthening process for the observed patterns, which leaves room for errors. Our data rules out the case of all-or-none memorization: Fig. S2 and Fig. S4 demonstrate that our subjects frequently made mistakes on specific patterns even after they were labeled correctly in previous presentations.

We can also rule out memorization of the types mentioned above as the sole mechanism. Relying on memorization alone would mean that all rules on patterns of three bits would be learned in the same rate. Fig S2 clearly shows that this is not the case, and that rule identity plays a key role. Even finer memorization aspects, such as pattern-specific acquisition rates, can also be ruled out from our data. Fig.S4 shows that subjects learned the labeling of the same pattern under different rules at different rates. Finally, pattern-response pairs can be influencing each other during learning. Fig. S3 complements Fig. S2. showing that the rule identity impacts this influence as well.

To hone in the general rule-based behavior we add Fig. S6 to show specific cases of pattern specific performance deterioration accompanying general performance increase. The examples in Fig. S6. Specifically highlight learning sessions in which a pattern-specific performance was high at the end of one session and decreased following a rule switch. Importantly, the rule switch did not require changing the learned response to the specific pattern (as 4 out of 8 patterns did not change response category in the rule switch). The pattern-specific performance deterioration during a general performance improvement is not expected in learning by stimulus-response association and is a hallmark of rule-based behavior.

### Behavior stability tests

#### Feature based behavior stability

We want to summarize how consistent were the monkeys in a single number for each session (Fig.S5A). This is done with answers from the last 1/4 of each session. We define as a consistency measure the mean (across patterns) distance of the logistic classifier (fitted to answers in the last 1/4 session) from the chance (0.5) answer.

Namely, if the monkeys adopt a feature based consistent policy at the last quarter of each session, then we can fit their sequence of answers with:

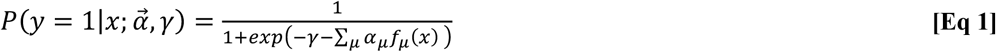

where *x* are the presented patterns, *f*_*I*_(*x*) are the features, and 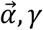, *γ* are fitted to maximize the likelihood of the answers.

A key reason to fit this classifier and not to make the consistency estimation per pattern is that there are much fewer pattern presentation per pattern and in any way calculating per pattern imposes the assumption that the monkeys can tell all patterns apart from each other. The classifier’s way doesn’t make any assumption beyond a features based behavior policy.

The consistency measure is thus the mean distance from 0.5. or,

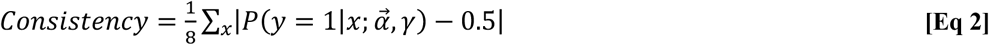

The chance level for a completely unbiased classifier is 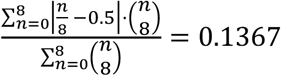

#### Across sessions stability

Next, we want to check if the monkeys were stable across sessions. Namely, regardless of performance, how similar is the features-based behavior at the last quarter of different sessions. Or, how similar is the behavior when learning the same rule in different sessions.

To avoid cross interference between the within-trial variability (the inconsistency that drives *P*(*y*|*x*) in Eq. 1 close to 0.5) and the across-trial variability (The inconsistency that separates the logistic classifiers that are fitted to different sessions) we threshold the classifiers, *y*(*x*) = [*P*(*y*|*x*) > 0.5], and for each rule compare all pairs of sessions. Fig.S5B shows the mean (and SE in errorbars) of the across session similarity score:

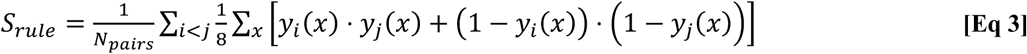

where *i, j* are different sessions of the same rule and *N*_*pairs*_ is the number of pairs of sessions with the same rule.

#### Behavior similarity across rules

To estimate how similar is the monkeys’ behavior to the rules they learn we repeat the calculation in Eq. 3 but replace one of the classifiers (*y*_j_) and subtract 0.5 to shift the mean expected overlap to 0. The, above chance level, results are presented in Fig.S5C.

#### Testing for performance-based strategy

Animal behavior could potentially obey a local performance-based strategy called win→stay, lose→switch. This strategy is observed in animal studies and suggests that animal will repeat a choice that led to reward and switch a choice that didn’t. To test if the monkeys significantly relied on such a strategy we simulated answer sequences following this strategy for all the learning sessions in our experiments. We than compared the true answers, given by the monkeys, to the simulated sequences. Any above-chance (50%) agreement would indicate that the monkey might be using this strategy. However, we find that the agreement with the win-stay, lose-switch strategy is below chance for nearly all sessions. Fig. S7 shows distributions of per-session agreement for simulations initiated in the left or right choice (the only free parameter). 93.5% of sessions had below-chance agreement and the median agreement was significantly lower than chance (Wilcoxon signed rank test. p<1e-15)

### Data Analysis – Neural activity

#### Single-neuron responses

We expect the learning-relevant cognitive mechanisms to be influenced by both trial-by-trial variations, such as changing behavior and stimulus identity, and by slower processes, namely learning. Studying the learning related dynamics, we are interested in the single unit neural activity that correlates to such inherently variable computational primitives. Namely, we seek a measure of the spiking activity that communicates the variations across trials. Accordingly, for every neuron we examine the spikes in the 500ms following the stimulus onset and bin them into 5×100ms segments to obtain sensitivity to temporal effects in addition to the spike count. The result is a 5-vector of spike counts from each trial, 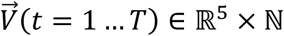. In this representation, the component of largest across-trials variance is 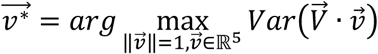. We then project each 5-dimensional vector on this principle component and get a single number from each trial, 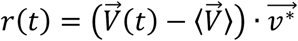. This number scores the spiking patterns of the neuron with respect to its most prominent fluctuations or change. Importantly, several unrelated processes may contribute to the across-trials variability and in choosing the projection, *r*(*t*), as the representation of stimulus neural response, we tune to the largest source of variability, regardless of its nature (Fig. S10A-C).

#### Pattern-specific neurons

We devised a criterion for exemplar preference based on a neuron’s firing rate in the 500mSec after the pattern presentation. We build a table of all responses of the neuron to each of the patterns. Next, we compare the sets of responses to each pair of patterns using a rank-sum test for equal medians and treat the distribution of responses as different using a threshold at p<0.05. A neuron is pattern-specific if the distribution of responses to only one pattern is different from all the others (Fig.S9)

#### Feature-based representation of the rule and neural responses

For a pattern, 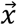, we chose a basis of features that are polynomials of the variables *x*_1_, *x*_2_, *x*_3_ that take the values ±1. Features, 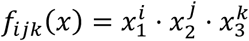 differ in the polynomial degrees (*i, j, k* = 0,1) and satisfy, when averaging across all patterns, ⟨*f*⟩ = 0, ⟨*f*^2^⟩ = 1.

##### Lemma 1

Different features in this base are statistically independent.

**Proof 1**: Let *f*_1_ and *f*_2_ be features in this set and without losing generality assume that they differ in the polynomial degree of *x*_1_ s.t. *f*_2_ doesn’t contain *x*_1_. Since ∀_*i*_, *P*(*x*_*i*_ = 1) = 0.5 we get that *P*(*f*_1_ = 1|*f*2) = *P*(*x*_1_ = 1) = 0.5 = *P*(*f*_1_ = 1).

##### Lemma 2

Correlation projections factor in this basis

**Proof 2**: Let ***f*** be a vector in features space, ***f*** = ∑_*I*∈{*i,j,k*}_ *a*_*I*_*f*_*I*_ such that 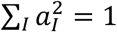. This means that ⟨***f***⟩ = 0 and ⟨***f***^2^⟩ = 1 (because ⟨*f*_*I*_ ⋅ *f*_*j*_⟩ = 0, ∀_*i*≠*j*_). Which leads to *Var*(***f***) = 1. So, if *n* is some random variable (say, the neural projection) the correlation 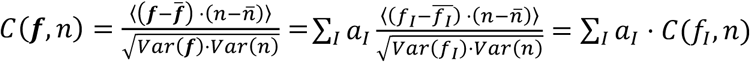.

This means that if we measure the correlations to the features separately, then the unit vector, *a*, that maximizes ∑_*I*_ *a*_*I*_*C*(*f*_*I*_, *n*) will give us the preferred feature. Also, if *a* is a rule that we chose in advance, e.g. the one being learned, then the projection ∑_*I*_ *a*_*I*_*C*(*f*_*I*_, *n*) is indeed the rule-correlation (Fig.3B, Fig. S10C,D).

#### Dynamics of representations

To study the dynamics of task related neural correlates we divided each session to partially overlapping windows (40 trials segments with 4 trials jumps). For each neuron, calculating the correlation between its spiking patterns, *r*(*t*), following stimulus onset, and the stimulus features, 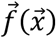, (as well as to the correct category and the monkey’s future answer) yields a set of correlation coefficients, 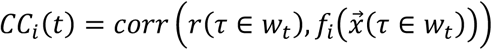, for each regression window *w*_*t*_. These rolling regression coefficients were used to calculate the following measures:

#### Comparing representation between conditions

To judge whether neurons show rule selectivity during a certain segment of the session (Fig.2D) we test the fraction of regression windows within that segment, that exhibit significant rule correlation (Pearson, p<0.05). This test is done comparatively between sessions of different conditions, and we set a criterion of 10% to declare a neuron as showing rule selectivity during the segment. If there are more than 5 neurons meeting each condition (2 conditions, e.g. easy and hard rules) we use the binomial comparison z statistic, 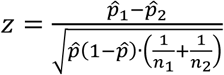 with 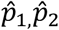 the measured success rate in two populations of sizes *n, n* and 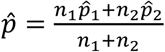.

#### Comparing rule vs. answer representations

Since these correlations have a mutual component (the spiking pattern) and interrelate via the performance level, we compare with William’s t-statistic for correlated correlations, 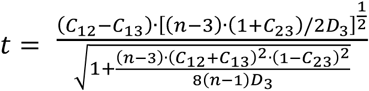, where *C*_12_ is the category correlation, *C*_13_ is the answer correlation, and *C*_23_ is the correlation between answers and categories. *n* is the number of trials and *D*_3_ is the determinant of the sample correlations matrix. The statistic is compared to the t – distribution with *n* − 3 degrees of freedom (Fig.2F).

#### Relating neural representation to behavioral performance

To relate any regression measure and performance within a group of neurons we take the following steps:

1. For every 40-trials-long regression window we calculate the mean performance.
2. Given a performance level, we collect all the regression windows with performance within 0.15 of that level and calculate the mean and standard error of the measures of interest (Fig. 2E).

#### Angle-to-rule and vector-magnitude

Given a basis of visual features there is a unique spanning of the classification rule in each session. For each regression window we define the angle-to-rule as the angle between the vector of correlation coefficients to visual features and the vector that represents the rule. Similarly, we define the features’ correlation magnitude as the norm (L2) of the correlation coefficients vector.

When presenting the learning related dependence of these geometrical variables over time, we smooth them with 10 percent of the running windows in a session (Fig.3, Fig.4D-G insets, Fig.S14- S16)

#### Correlation to rule and answer

In Fig. S14 and Fig. S16. we present similar analyses of representation dynamics (as in Fig. 4D-G) but instead of the geometric measures we show correlations to the categories determined by the rule (annotated as ‘C’ in Fig. S14) and correlations with the monkeys physical answers (their right or left choices in each trial, annotated by ‘A’ in Fig. S16)

#### Session-length standardization

Several calculations require the comparison or grouping of segments from relative session fractions and/or location. To enable this, we standardized the regression measures from each session to a fixed length of 100 bins. This means that all rolling regressions were stretched to the same length, because a hundred regression windows would only come from 436 trials in a session.

#### Cells that reduce angle-to-rule or increase vector-magnitude

To quantify neurons that decreased angle-to-rule or increased the vector-magnitude (Fig.4D-G), we compare regression windows in the first 15%-segment of the sessions to regression windows in the last 15%-segment of each session with a 1-tailed t-test. The fractions of cells that passed the test are compared with a 2-tailed binomial z test.

#### Fractional change in neural-vector, angle and magnitude

We calculate the fractional difference from their average baseline values in the initial fraction of regression windows (Fig.4). The resulting traces are smoothed with a 10% window and significant difference between sessions of easy and hard rules is determined with bootstrapping – shuffling the easy/hard label 10,000 times and checking if the correct labeling surpasses the required confidence level (95%).

#### Optimal lags between time series

Given two time series, e.g. the angle-to-rule of a dACC neuron and the simultaneously-recorded vector-magnitude of a Putamen neuron, we find the shift that maximizes their Pearson correlation. Only pairs with significant correlation in the optimal lag contribute (as in Fig.4I,J).

#### Relating the neural-vector to next-day behavior

To examine if the learning-related change in the neural-vectors indicate a real shift in the monkeys’ preferred policy (Fig.4K,L), we tested if the neurons’ preferred feature combination (i.e. their neural- vector) predicts the monkeys’ behavior early in the following day. For each neuron we averaged the neural-vector in the late fraction of the rolling regression windows. Then, as a measure of similarity, we calculated the projection of the neural-vector on the subsequent day’s rule. In Fig.4K we calculate the Pearson correlations between these neural projections and the mean performance in the early fraction of the next day’s session across the neural population. In Fig.4L we repeat the same calculation but only take cases in which the rule was changed between the current and next day.

#### Code and Data availability

Custom code for behavioral and electrophysiological tests is available from the corresponding author upon reasonable request.

All data supporting the findings of this study are available from the corresponding author upon reasonable request.

