## Supplementary Figures for "A geometric representation unveils learning dynamics in primate neurons"

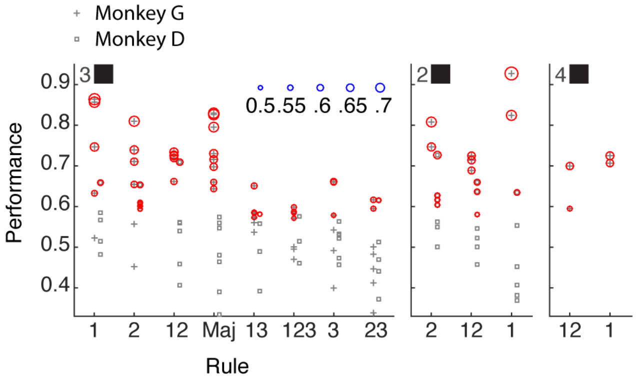


**Fig.S1. Performance in all session and rules.** Identical to Fig. 1D but with ordering the rules according to mean performance (left to right). Gray markers show performance at the last quarter of sessions ($P_{end}$, see methods) and, for significant results, the red circles' size shows the confidence level. Both animals had atleast one successful session in each rule, but four rules (left) had many more successful sessions than the other four (right).

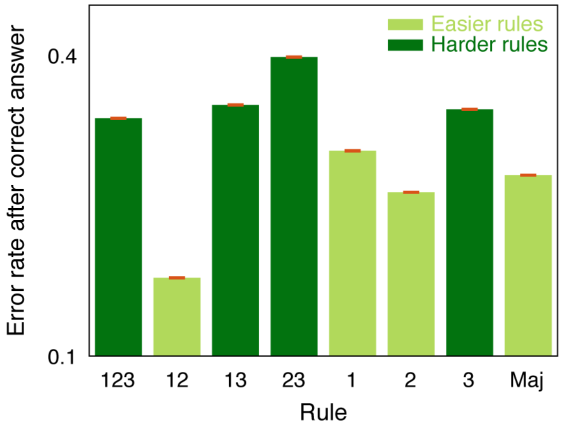


**Fig.S2. Error probability after making a correct classification.** For each rule (x-axis) we plot the probability of making an error (y-axis) after choosing correctly in the previous presentation of the same pattern. Error bars depict SE (red). The relatively low probability and the strong dependence on the specific rule suggests the monkeys did not memorize specific patterns after a successful classification. Kruskal-Wallis test across all repetitions and patterns validates the effect of the rule on the occurrence of errors after correct classifications ($\chi^2(7)=88.97$, $p < 1e\text{-}15$). Pairwise binomial z-test shows that all the error rates in easier rules are smaller than error rates in harder rules.

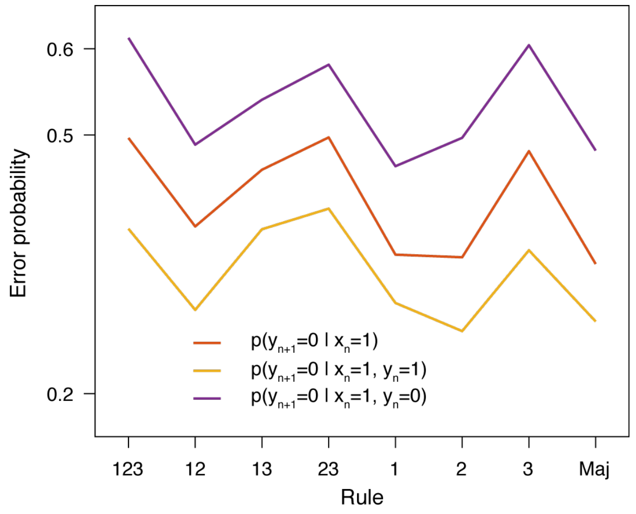


**Fig.S3. Error probability in different stimulus-response associations show rule-dependent coupling.** For each rule (x-axis) we plot the probability of making an error in classifying a pattern 'y' (y-axis) after correctly-classifying a different pattern, 'x', in the previous presentation of 'x' (red line). In yellow (purple) we plot the same error probability conditioned also on correctly (incorrectly) classifying 'y' in the previous presentation. The latter calculation shows the global effect of learning – fewer errors after correct choices. All probabilities have strong dependency on the rule being learned (Kruskal-Wallis test, $\chi^2 > 88$, p<1e-10 in all cases) – showing the coupling between stimulus-response pairs expected in rule-based learning.

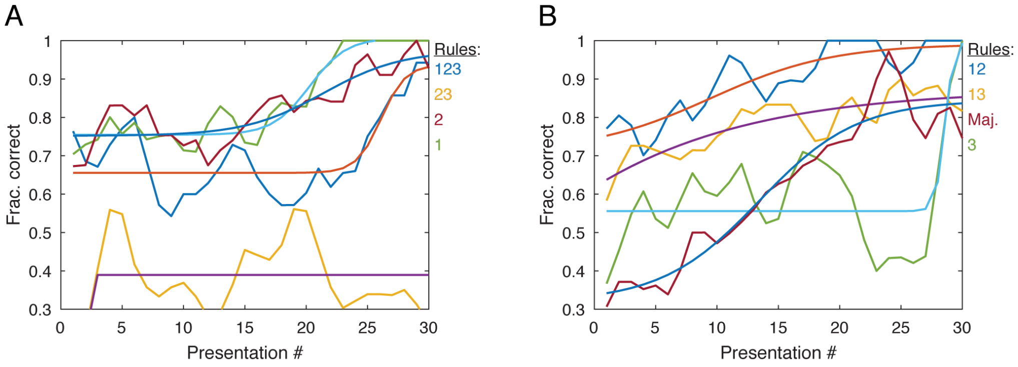


**Fig.S4. Acquisition rates for the 'salient' pattern.** The most salient pattern, '000', was labeled 'right' and 'left' equally in our set of rules (4+4 rules). Shown are the learning curves (colored lines + sigmoidal fits), the proportion of correct labeling (y-axis, smoothed across days with a running window of 4 presentations) along the pattern presentations (x-axis). The two panels are for the 4 rules in which 000 is labeled 'left' (**A**) and for the 4 it was labeled 'right' (**B**). There was variability across rules in how this most visually salient pattern was learned, showing that patterns that draw more attention are not learned faster or similarly across different rules, and therefore suggesting against simple memorization process. Kruskal-Wallis test across all segments of 10 pattern presentations validates the effect of the rule on the performance in panels a,b. ($24 < \chi^2(3{,}36) < 33$, 1e-7 < p < 1e-4 in all segments).

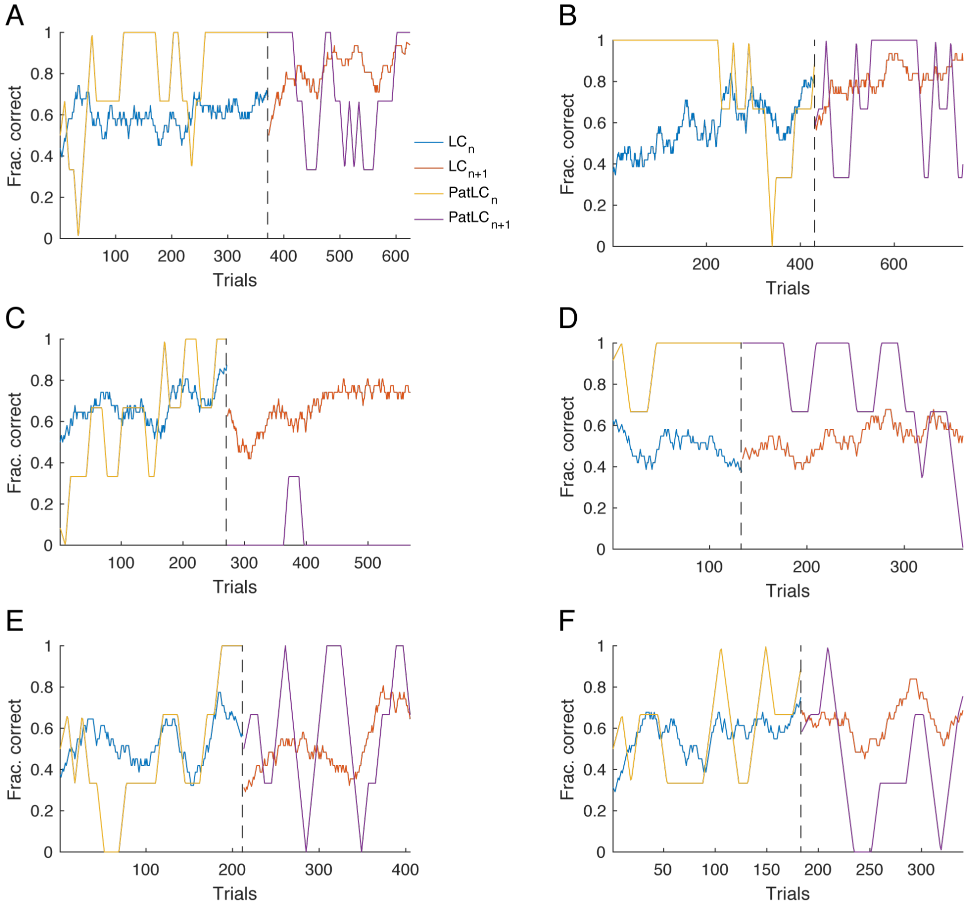


**Fig.S6. Learning a new rule disrupts acquired stimulus-response associations that agree with the new rule.** Blue and red lines show general performance in consecutive sessions. The rule determining the correct answers was switched between the sessions (dashed line) but 4 out of 8 patterns did not change their associations. The yellow and purple lines show the performance in such patterns. In contrast to learning by independent stimulus-response pairing, rule-based learning predicts that pattern-specific performance can deteriorate while the general performance improves. (**A-D** for monkey G. **E,F** for monkey D). This can happen even if the association-specific performance reaches 100% (**A,E**).

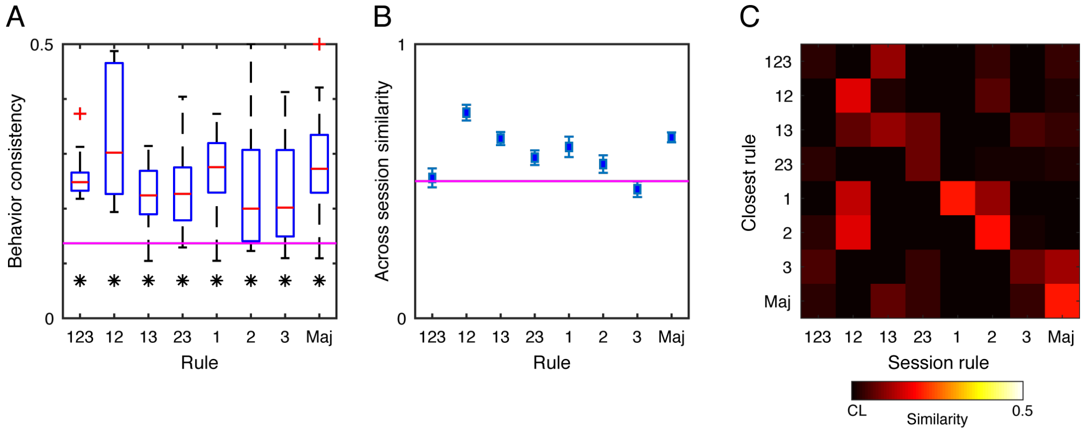


**Fig.S5. Behavior consistency. A.** The within-session consistency scores (y-axis, methods) for all sessions with a single rule (x-axis). The chance level is marked by the magenta line. Statistical significance (t-test, $p<0.05$) of above chance mean is marked by asterisks. **B.** The between-session pairwise consistency score (y-axis, SE in error bars) across sessions with a single rule (x-axis). The chance level is marked by the magenta line. **C.** The classifier, fitted to the last ¼ of each session in which a certain rule was taught (x-axis), is compared to the set of rules (y-axis, methods). The color scale is from the chance level (CL) to the maximal possible value (0.5). Please see methods for full description of the measures

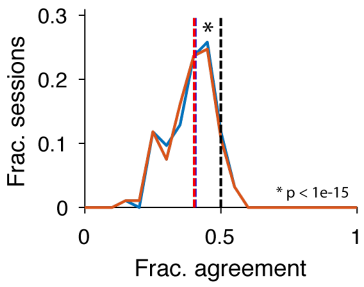


**Fig.S7. The Win→Stay, Lose→Switch strategy poorly describes behavior.** We created simulated win→stay, lose→switch answer sequences, using the same pattern presentations and rules as the true sessions. The curves show histograms of the overlap fraction between the simulated sessions with the true behavior. (Red and blue curves are for simulations starting with the left and right choices). Colored dashed lines show the median values, both smaller than the chance level (0.5, black dashed line, Wilcoxon's signed ranked test, p<1e-15).

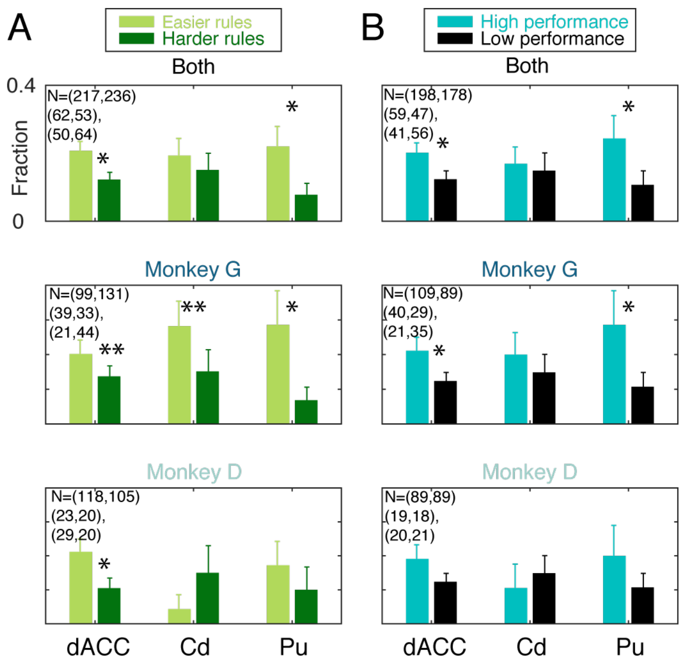


**Fig.S8. Robustness of rule-representation in single-units**. Similar presentation to Fig.2d: The fractions of neurons that exhibited significant rule correlation during the last third of the session (bars, y-axis. SE in error-bars) are plotted for the 3 regions (x-axis). Apart from grouping both monkeys (top) we also separate to neurons from monkey G (middle) and D (bottom). **a.** Separating sessions by the rule-group. Easy rules in light green and hard rules in dark green (Both: dACC: $z=2.43$, $p<0.01$, Pu: $z=2.12$, $p<0.02$; Monkey G: dACC: $z=1.305$, $p<0.1$, Cd: $z=1.3275$, $p<0.1$, Pu: $z=2.37$, $p<0.01$; Monkey D: dACC: $z=2.17$, $p<0.02$). **b.** Separating sessions by performance in each session separately (i.e. independent of the rule). (Both: dACC: $z=2.046$, $p<0.021$, Pu: $z=1.79$, $p<0.04$; Monkey G: dACC: $z=1.85$, $p<0.04$, Pu: $z=1.97$, $p<0.03$**).**

Pairs of N values show number of neurons in dACC,Cd,Pu and each condition (Easier,Harder) or (High performance, Low performance). Significant differences by binomial z-test (*=p<0.05, **=p<0.1).

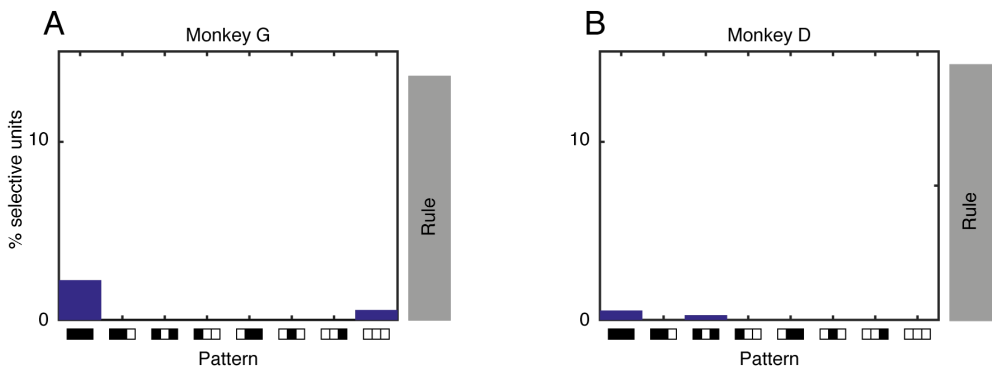


**Fig.S9. Pattern selective neurons.** The percent of neurons that were selective for a single pattern (bars, y-axis) is plotted for all patterns (x-axis). **A.** monkey G. **B.** monkey D. Gray bars indicate the percent of rule correlated neurons.

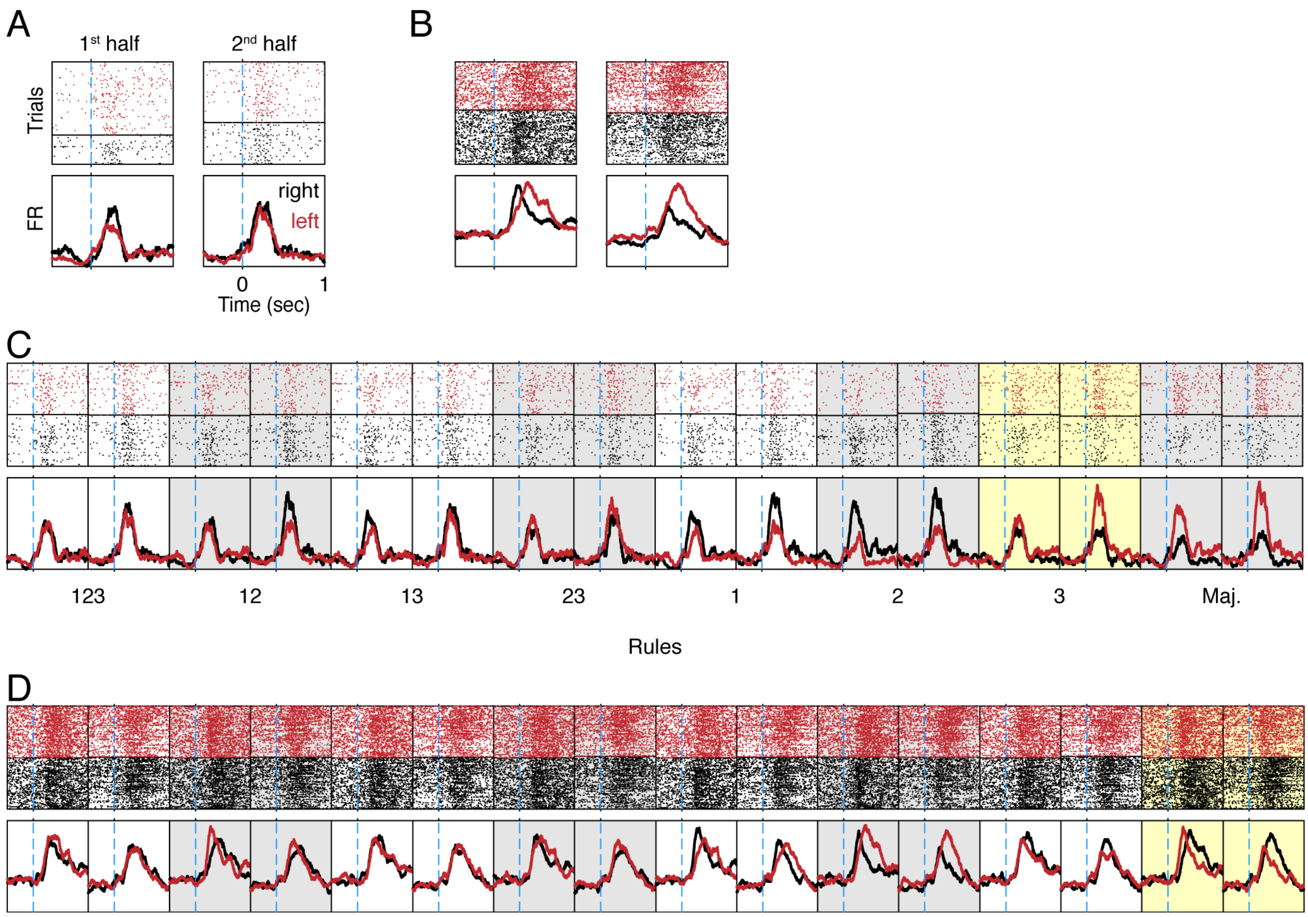


**Fig.S11. Spike rasters and PSTHs calculated in the 1st and 2nd halves of learning sessions show single neuron changes in representing the animals' answers, correct rules, and other rules. A.** Spikes (of one neuron ,left in Fig. 2c) are aligned to the time of stimulus onset (t=0, x-axis, light blue dashed line). Trials (y-axis) are stacked according to the animal left or right choice (red, black dots in the top panel and curves in the bottom panel mark spike times and firing rates accordingly). Calculations are made separately in the first and second halves of the session (left, right panels) to show that this neuron doesn't differentiate the animal's choice in the 2nd half of the session. **B.** Like panel A but for a different neuron. In this case the PSTHs are different for the different choices in both parts of the session. **C.** The same data as in panel A separated by the labels of the 8 rules in our experiment (x-axis labels). Gray and white shadings distinguish the different rules and yellow shading marked the actual rule being learned. **D.** similar to panel C but for the data in panel B.

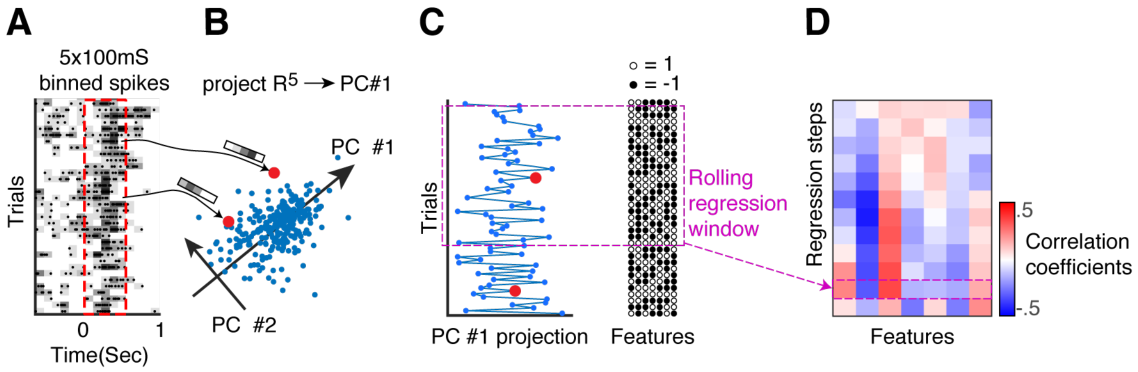


**Fig.S10. A schematic showing the calculation of single neurons' dynamic feature representation.** **A.** Spikes (dots) during 500 msec after the stimulus presentation (x-axis, dashed red frame) are counted in 5 x 100 msec bins (gray scale). **B.** The 5-vectors of spike counts from all trials are projected on the principle component explaining the most variance across trials. (arrows and red dots show specific examples) **C.** The resulting neural 1-d stimulus response, x-axis, is correlated across trials (y-axis) with stimulus visual features (7-vectors of black and white circles) in a rolling regression window (purple frame). **D.** The resulting vector of 7 correlation coefficients spans the neuron's visual feature preference during that regression window and its dynamics across regression steps (y-axis)

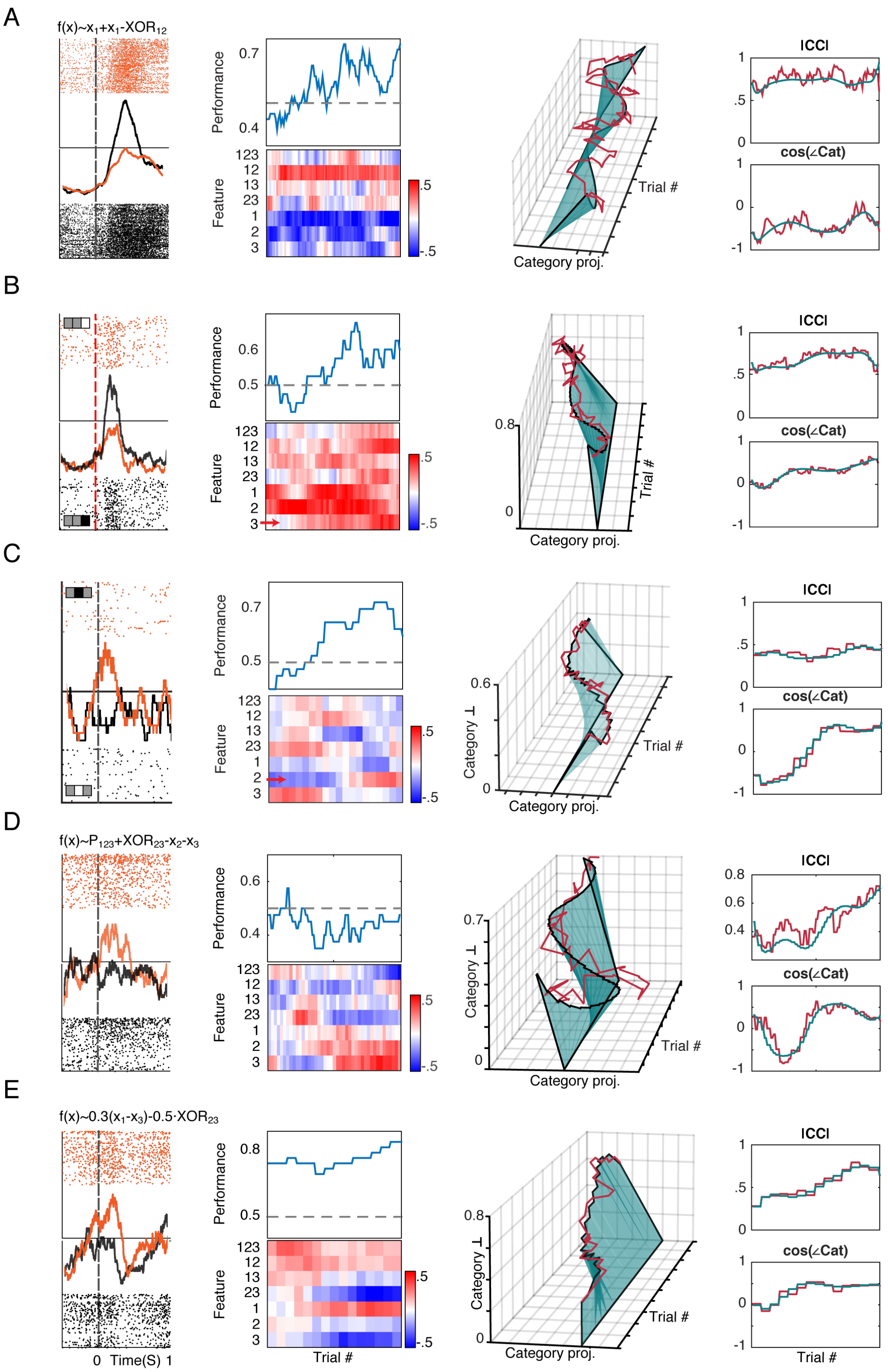


**Fig.S12. Examples of learning related neural dynamics.** Same format as in Fig.3, showing neurons with stable feature selectivity, high dimensional rotation, magnitude stretching, and complex trajectories.

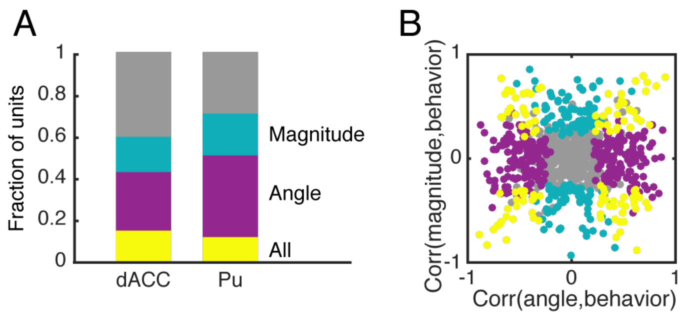


**Fig.S13. Significant correlations between geometric neural correlates and learning curves partially overlap. a.** Enumeration of the significant (Pearson, $p<0.05$) correlations to all variable combinations in the different regions (x-axis). The colored bars show the fraction of units with significant correlations between the learning curve and both angle-to-rule and correlations vector-magnitudes (yellow), only angle-to-rule (purple), or only vector-magnitudes (turquoise). **b.** For each neuron we compare the magnitude correlation (y-axis) to the angle-to-rule correlation (x-axis). Color-coding as in panel (a).

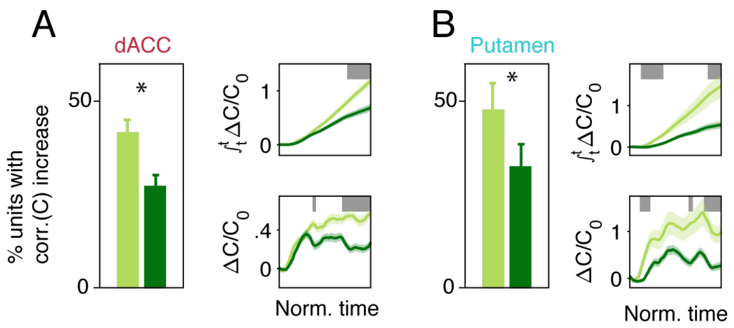


**Fig.S14. Dynamics of category correlations are similar to angle-to-category.** Panels A,B repeat panels D,E in Fig. 4 but for the value of category-correlation coefficients – showing very similar effects.

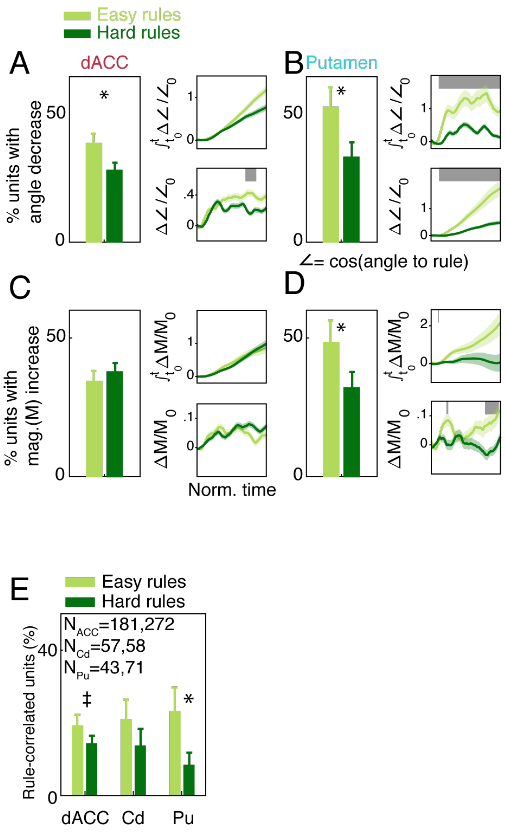


**Fig.S15. Similar results when redefining the Majority rule as hard for monkey D. A-D** repeat panels D-G in Fig. 4. With the same results. **E.** Repeats panel D in Fig 2. The binomial z-test comparing population sizes keeps the same trend (‡: $p < 0.1$, *: $p < 0.05$)

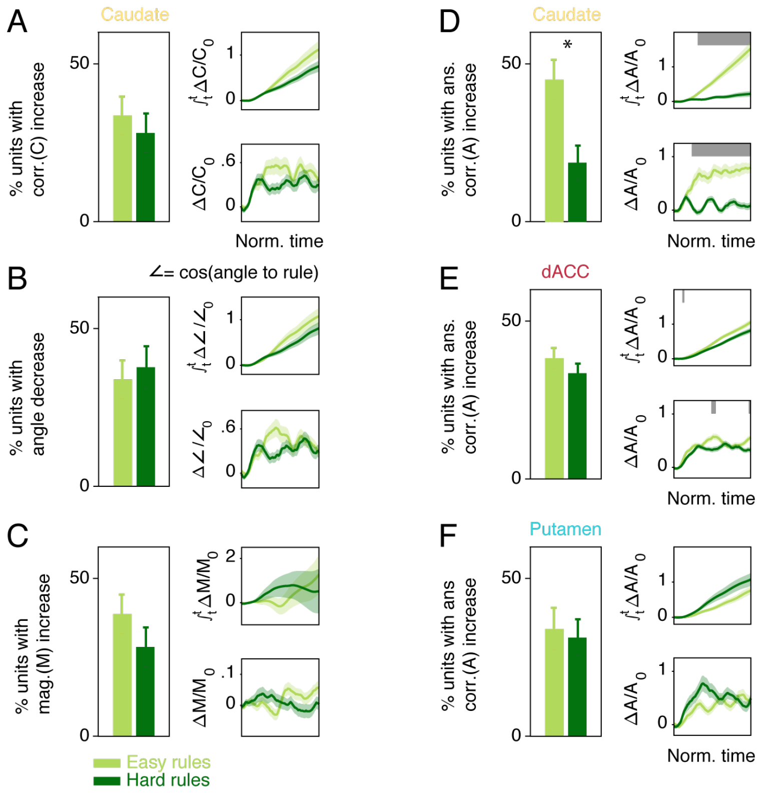


**Fig.S16. Neural representation in the Caudate. A.** Caudate neurons show a rule-dependent increase in representing the animals' actual-choice (Fig. 2F) but not the correct label. **B,C.** Repeat Fig. 4D,F for neurons recorded in the Caudate. Panels A-C show that the caudate population doesn't move to represent a learned rule. **D.** Proportion of neurons that significantly increased their correlation to the animals' choices (A – correlation to answers) during the session was different in easy rules vs. hard rules in the Caudate ($p<0.05$, binomial z-test). Right panels show the average change in correlation coefficient (bottom, SE in shaded color) and the normalized cumulative change (top). Sessions were time-warped for averaging. Gray bars indicate significant difference between easy and hard rules ($p<0.05$, bootstrap). **E,F.** In contrast to the Caudate, the dACC (E) and Putamen (F) do not show a differential change in representing the animals' choice in easier rules.

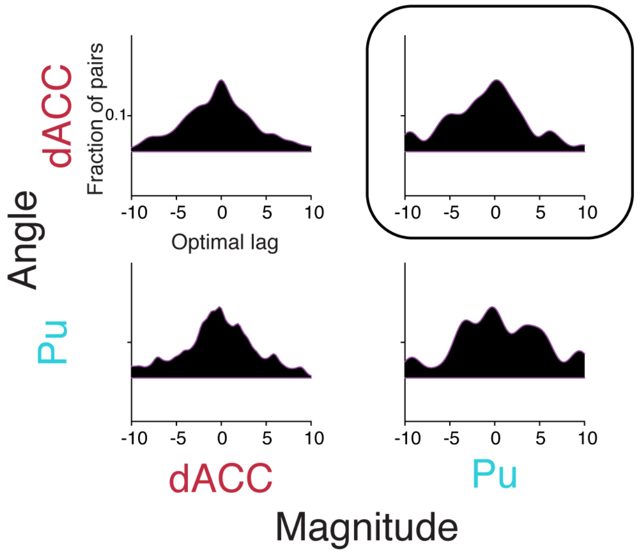


**Fig.S17. Optimal lags between neural-vector properties (magnitude and angle) between dACC and Striatal pairs.** For all simultaneously recorded pairs, we computed the optimal lag between the vector-magnitude and the angle-to-rule, for all four possible combinations. Only lags between vector-magnitude in Putamen neurons and angle-to-rule in dACC neurons were significantly different than zero (top-right, also shown in main Fig.4i,j) with the Putamen magnitude following dACC rotation ($p<0.004$, t-test; Mean lag $-0.7\pm0.24$ regression steps; all other comparisons were not different than zero, $p>0.1$ for all, t-tests).
